# Long term outcomes of Everolimus therapy in *de novo* liver transplantation: a systematic review and meta-analysis of randomized controlled trials

**DOI:** 10.1101/703215

**Authors:** Bzeizi Khalid, Smith Richard, Benmousa Ali, Dama Madhukar M.V.SC, Aba-Alkhail Faisal, Jalan Rajiv, Broering Dieter

**Affiliations:** Liver Transplant Department, King Faisal Specialist Hospital and Research Center, P.O. BOX 3354, Riyadh, 11211, Saudi Arabia; Department of Medicine, Ipswich Hospital Trust, Heath Rd, Ipswich IP4 5PD, UK; Perfect Statistics, Hubli, India 580023; Multi Organ Transplant at King Fahad Specialist Hospital, Dammam, Saudi Arabia.; Liver Failure Group, UCL Institute for Liver and Digestive Health, UCL Medical School, Royal Free Campus, London NW3 2PF, United Kingdom.; Department of Liver Transplantation and Hepatobiliary Surgery, King Faisal Specialist Hospital and Research Center, P.O. BOX 3354, Riyadh, 11211, Saudi Arabia.

**Keywords:** Calcineurin inhibitor, everolimus, liver transplantation, long-term, withdrawal

## Abstract

**Background:** Risk of nephrotoxicity in liver transplant patients on calcineurin inhibitors (CnIs) is a concern. Several controlled trials reported benefit of Everolimus (EVR) in minimizing this risk when combined with a reduced CnIs dose.

**Objective:** To systematically review the efficacy and safety of EVR, alone or with reduced CnI dose, as compared to CnI alone post liver transplantation.

**Methods:** We searched MEDLINE, Scopus and the Cochrane Library for randomized controlled trials (RCTs) comparing EVR and CnI based regimens post liver transplanation. Assessment of studies and data extraction was undertaken independently.

**Results:** Eight studies were selected describing 769 patients. Cockcroft-Gault GFR (CG-GFR) was significantly higher at one (p=0.05), 3 & 5 years (p=0.030) in patients receiving EVR as compared to those receiving CnI therapy. The composite end point of efficacy failure was similar between the two arms after 1, 3 & 5 years of study. Higher number of patients discontinued EVR due to adverse effects in one year, however no difference was noted after 3 & 5 years. A higher rates of proteinuria, peripheral edema and incisional hernia were noted in patients on EVR.

**Conclusion:** The analysis confirms non-inferiority of EVR and reduced CnI combination. Patients on the combination regimen had better renal function compared to standard CnI therapy.

## INTRODUCTION

Chronic renal dysfunction is an important cause of mortality and morbidity following liver transplantation.^1^ Although, the indications, techniques, patient selection, and immunosuppressive therapy used for liver transplantation have evolved, renal dysfunction remains as an important limiting factor.Approximately18% of patients develop chronic renal failure or end stage kidney disease by five years post-transplant.^2^ Various factors such as pre-transplant renal status, female gender, age, presence of hepatitis C virus (HCV) infection and calcineurin inhibitors (CnI) therapy influence the deterioration in renal function.^2^

Calcineurin inhibitors, the cornerstone of immunosuppression post liver transplantation, are an important modifiable risk factor for renal dysfunction.^3^ Several clinical trials have investigated the risk associated with use of CnI therapy and how the deterioration in renal function can be ameliorated.^4,5,6^ The comparators for such evaluations are the mammalian target of Rapamycin (mTOR) inhibitors; sirolimus and everolimus (EVR).Everolimus gained approval for use in liver transplant patients following its introduction as an immunosuppressant in renal transplantation.^7^ Use of EVR is approved in combination with reduced dose tacrolimus (RTAC) after 30 days of liver transplant.^7^

Several studies have looked at the efficacy and safety of either EVR monotherapy or reduced CnI dose combination therapy (EVR+RTAC) compared to the standard CnI therapy post-liver transplantation.^8,9,10,11,1,12^ There have been significant differences in the study designs in the limited number of studies conducted so far. The multicentric H2304 study reported the results of comparison of EVR+RTAC with tacrolimus (TAC) control after one, two and three years of institution of therapy among *de novo* liver transplant patients.^11,12,13^ The PROTECT (Preservation of Renal function in liver Transplant rEcipients with Certican Therapy) trial, evaluated EVR monotherapy as compared to standard CnI therapy after one, three and five years after *de novo* liver transplant.^14^ Other single center studies have evaluated EVR monotherapy compared to EVR in combination with other immunosuppressive agents.^15^

These clinical studies showed non-inferior rejection rates with EVR (in either of the regimens) and less deterioration of renal function as compared to standard CnI therapy.^11,1,16,13^ However, new evidence regarding the incidence of adverse events (AE) with EVR has led to changes in the prescribing information of EVR. A recent US FDA update has recommended changes in prescribing information of EVR in cases of interstitial lung disease, non-infectious pneumonitis and pulmonary hypertension (including pulmonary arterial hypertension). Additionally, the clinical trials have shown an increased incidence of discontinuation of study treatment in the EVR treatment arms.^11,12^

There is a need for more evidence on both the long and short-term safety and immunosuppressive efficacy of EVR alone, or in combination with RTAC, as compared to standard CnI monotherapy. This systematic review addresses the efficacy and safety of EVR post liver transplantation.

## METHODS

- **This review has included** Randomized controlled clinical trials on *de novo* liver transplantation patients who received EVR as part of their immunosuppressive regimens in comparison to CnI based immunosuppression. EVR+RTAC (Reduced exposure tacrolimus) or EVR monotherapy was compared to the standard therapy with ≥6 months of follow-up. The following comparisons were included: EVR monotherapy versus standard CnI therapy, Addition of EVR versus placebo and EVR + RTAC versus standard CnI therapy

### Types of outcome measures

The outcomes or interest were change in renal function assessed by eGFR, treated biopsy proven acute rejection (tBPAR), graft loss, mortality, treatment emergent adverse events (TEAE) leading to withdrawal from therapy and hepatic artery thrombosis (HAT).

tBPAR had been defined as acute rejection with a locally confirmed rejection activity index (RAI) ≥3 according to Banff criteria treated with anti-rejection therapy.^11^

#### Search methods for identification of studies

### Literature search

Preferred Reporting Items for Systematic reviews and Meta-Analyses (PRISMA) guidelines were adopted for this systematic review^17^. Literature search from the earliest available date to 1^st^ of May 2017 was performed in PubMed/MEDLINE, Scopus, and the Cochrane Library databases using the keywords “everolimus” and “liver transplant” or “liver transplantation” or “hepatic transplantation” or “hepatic graft” or “LT.” Relevant clinical studies (unpublished and ongoing trials) were also identified in the ClinicalTrials.gov registry of clinical trials (http://clinicaltrials.gov/). The literature search was not restricted by language or year and included unpublished studies. The reference lists of included studies were also screened manually for additional studies. Trials published solely in abstract form were, however, excluded because the methods and results could not be fully analyzed.

### Data collection and analysis

All abstracts and titles were scanned by KB and RS independently. All potentially relevant articles were reviewed as full text. Any differences in opinion about the selection of articles were resolved by a third party.

#### Data extraction and Risk of Bias Assessment

KB and RS independently retrieved relevant patient and intervention details using standardized data extraction forms. Authors undertook all stages of study selection and data extraction independently. The risk of bias of eligible RCTs was assessed with the Cochrane collaboration tool.^18^ Disagreements between reviewers, if any, were resolved by discussion to obtain a consensus.

### Data analysis

Dichotomous data were expressed as Odds Ratio (OR) with 95% confidence intervals (CI). Cochran ‘Q’ and I^2^ statistics were used to assess the heterogeneity among the studies. The level of heterogeneity demonstrated by the I^2^ score was characterized according to standard guidelines as complete absence (0%), low (25%), moderate (50%), and high (75%) level. Fixed effect model was used for meta-analysis of variables with homogenous data with statistically insignificant heterogeneity. Random effects model was used for meta-analysis of variables with statistically significant heterogeneity. Effect size was measured using odds ratio with 95% confidence intervals. Robustness of the results was reconfirmed by conducting sensitivity analysis to understand if any study had a major influence on the combined effect size. The combined effect sizes were interpreted with due consideration for publication bias analyzed through bias plots. A two-tailed P value of less than 0.05 was considered statistically significant. All the meta-analyses and associated tests were performed in Comprehensive Meta-analysis (CMA) software, Version 2.

## RESULTS

### Study selection and description of included studies

Eight RCTs passed the inclusion criteria (Figure 1). The characteristics of the included studies were summarized in Table 1. It must be noted that we have treated each data point within study as separate entry for the meta-analysis. A difference in treatment schedule, dose of EVR or the follow-up duration was considered as a criterion for considering the data points separate. A total of 2189 patients randomized to treatment group and 2248 patients randomized to control group. These studies compared EVR alone or in combination with RTAC to standard therapy or placebo. From the 8 studies, four data points were available on EVR with CnI reduction therapy and 6 data points were available on EVR with CnI elimination therapy. One study initiated the therapy on day one of the liver transplantation and the rest initiated the EVR therapy on 30^th^ day of the transplantation. All studies except Masetti were multicenter international studies.

**Figure 1.**
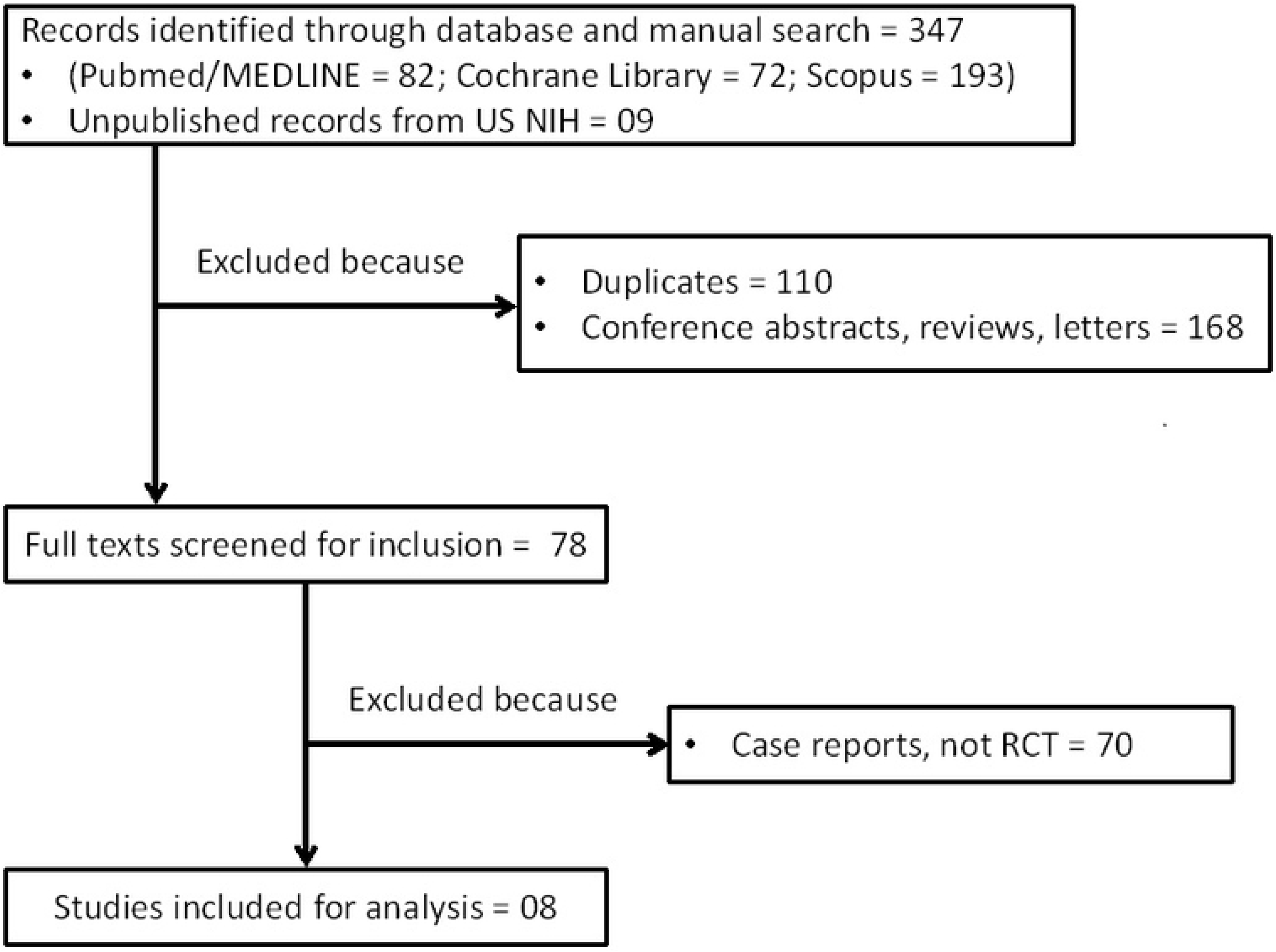

### Risk of bias

Included studies showed moderate risk of bias as assessed by the six items of the Cochrane instrument (Supplementary figures 1 and 2). All trials mentioned the method of randomization, but Levy et al, 2006^8^ did not specify the method of allocation. All RCTs except Levy et al, 2006^8^ were conducted with open-label design.

### Changes in renal function

In EVR + CnI elimination trials, the eGFR was significantly higher in treatment group (p<0.05) as compared to the control group (Figure 2a). The mean difference in treated patients was 20.33 mL/min, 14.57 mL/min, 9.47 mL/min, 16.30 mL/min and 11.70 mL/min at 6, 12, 24, 36 and 60 months respectively. In EVR + CnI reduction trials, the eGFR was significantly higher (p<0.001) as compared to the control group (Figure 2b), except at 12 months. The mean difference in treated patients was 8.55 mL/min, 6.90 mL/min, and 15.20 mL/min at 6, 24 and 36 months respectively. At 12 months, though the treated group had 3.73 mL/min higher value of eGFR than controls, this difference was not statistically significant.

**Figure 2.**
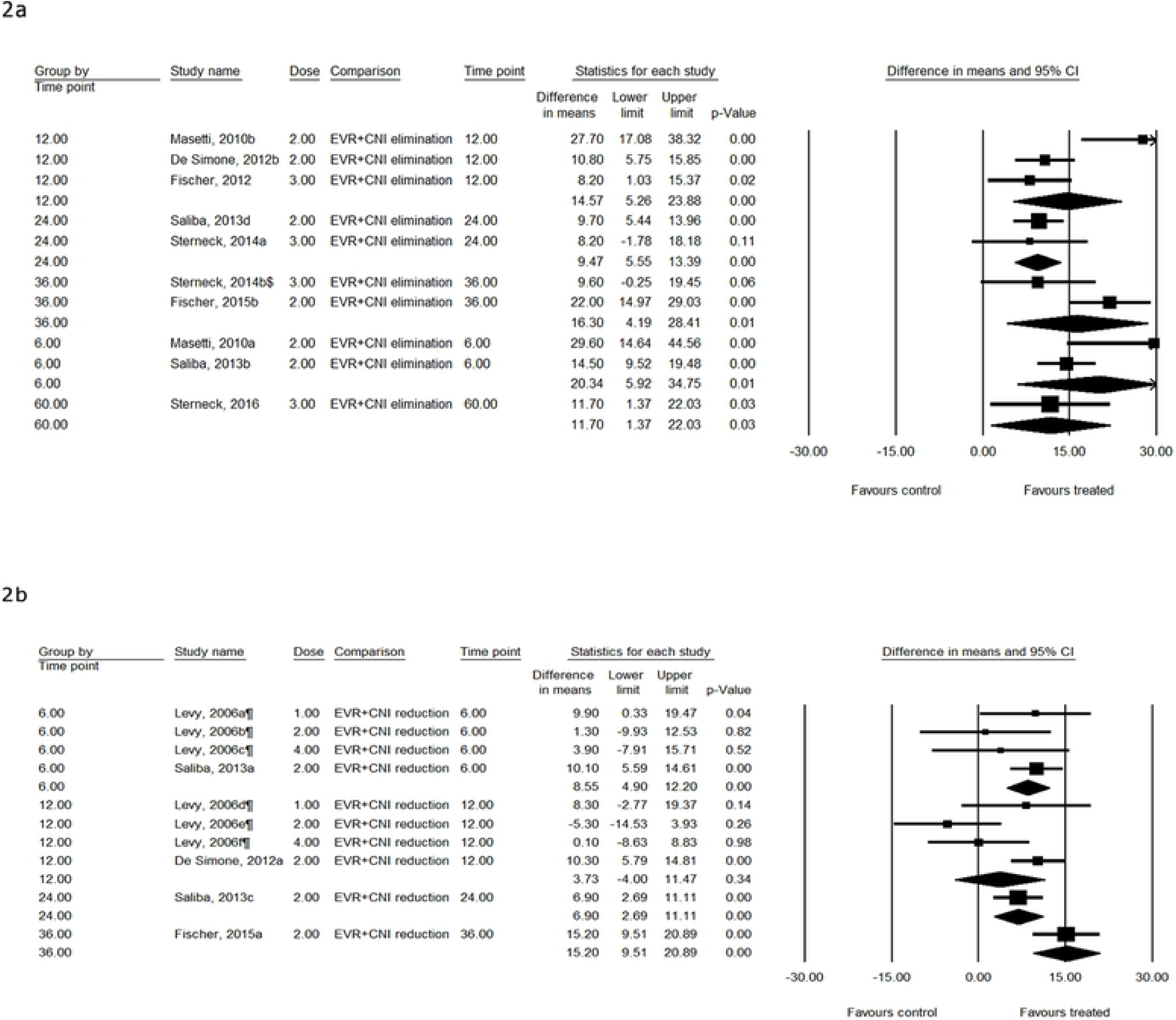

Subgroup analysis by dose showed that patients receiving EVR + CnI elimination therapy experienced an improvement of 17.03 mL/min and 9.16 mL/min in eGFR at EVR dose of 2 mg and 3 mg respectively. Similarly, patients receiving EVR + CnI reduction therapy showed a significant increase in eGFR at 1 mg (mean difference of 9.22 mL/min) and 2 mg (7.71 mL/min) dose of EVR (p<0.01). However, at 4 mg dose the difference was 1.44 mL/min (p>0.05).

### Treated biopsy proven acute rejection

The odds of tBPAR were significantly higher in patients receiving EVR + CNI elimination therapy (p<0.05). Patients in treatment group had 1.59, 2.06, 1.87 and 12.58 times higher odds of suffering tBPAR at 12, 24, 36 and 60 months post liver transplant (Figure 3a). On the contrast, the odds of tBPAR were significantly less in patients receiving EVR + CnI reduction therapy (p<0.01). Patients in treatment group had 0.48, 0.43, and 0.40 times lower odds of suffering tBPAR at 12, 24 and 36 months respectively after liver transplant (Figure 3b).

**Figure 3.**
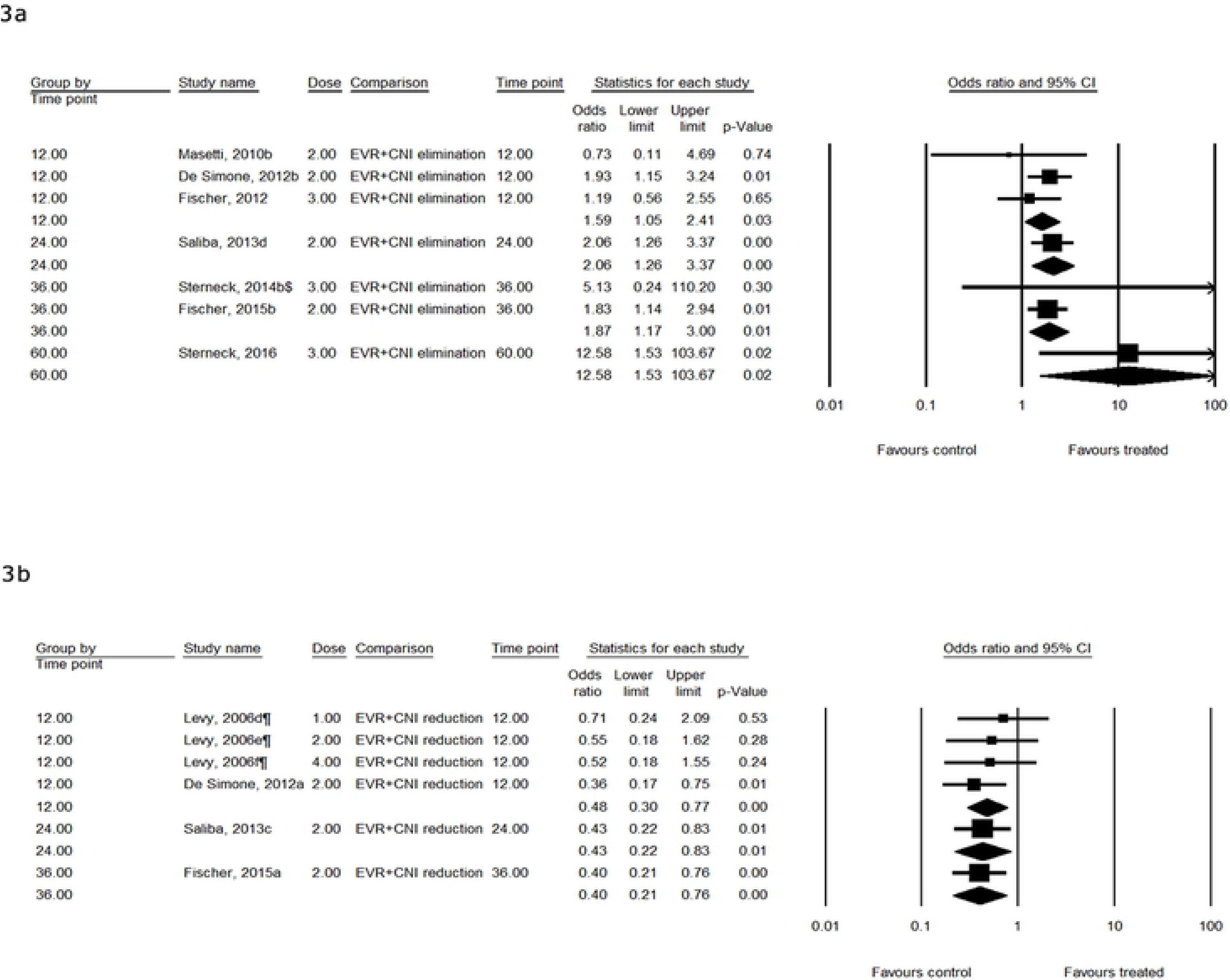

Subgroup analysis by dose of EVR showed that the odds of tBPAR were significantly higher in patients receiving a 2 mg of EVR in EVR + CNI elimination therapy group (p<0.05). Though the patients receiving 3 mg of EVR had an odds ratio of 3.18, the difference was statistically insignificant. In the trials with EVR + CnI reduction therapy, tBPAR was significantly less in treated patients at a dose of 2 mg (OR=0.48; p=0.00). However, at doses 1 mg and 4 mg, there was no difference between treatment and control groups.

### Graft loss

Graft loss rates were similar (P>0.05) between treatment and control groups of both therapy schedules (Figure 4a and 4b), for all the doses and at all the time points after liver transplant.

**Figure 4.**
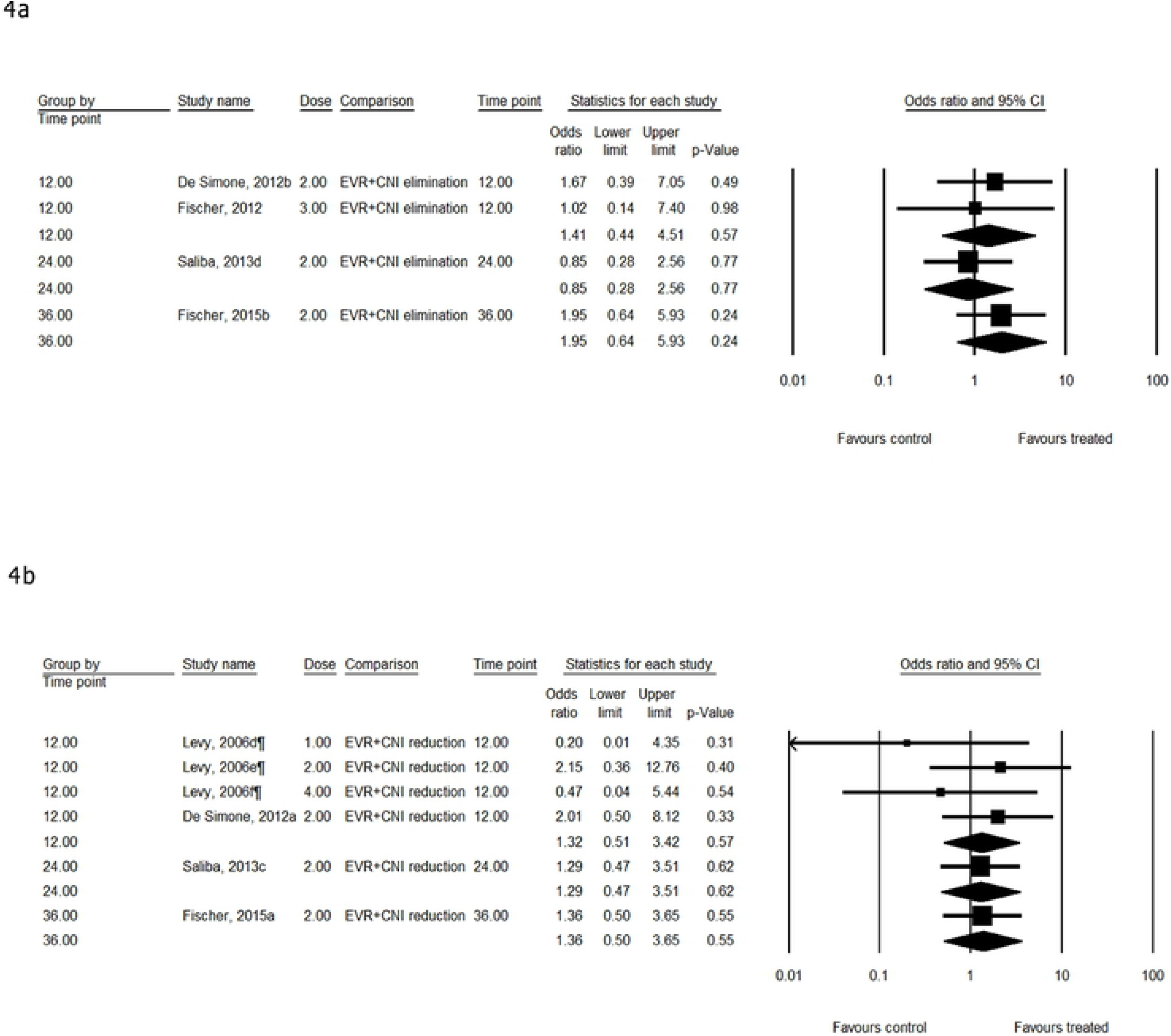

### Mortality

Mortality rates were similar between treatment and control groups of both the schedules (Figure 5a and 5b) at all the time points post liver transplant. In the EVR + CnI elimination trials, 2 mg dose of EVR was associated with significantly higher mortality rate as compared to control group (OR=2.06; p=0.04). However, mortality in patients receiving 4 mg was similar to control group. In the reduction group, the dose had no effect on the mortality.

**Figure 5.**
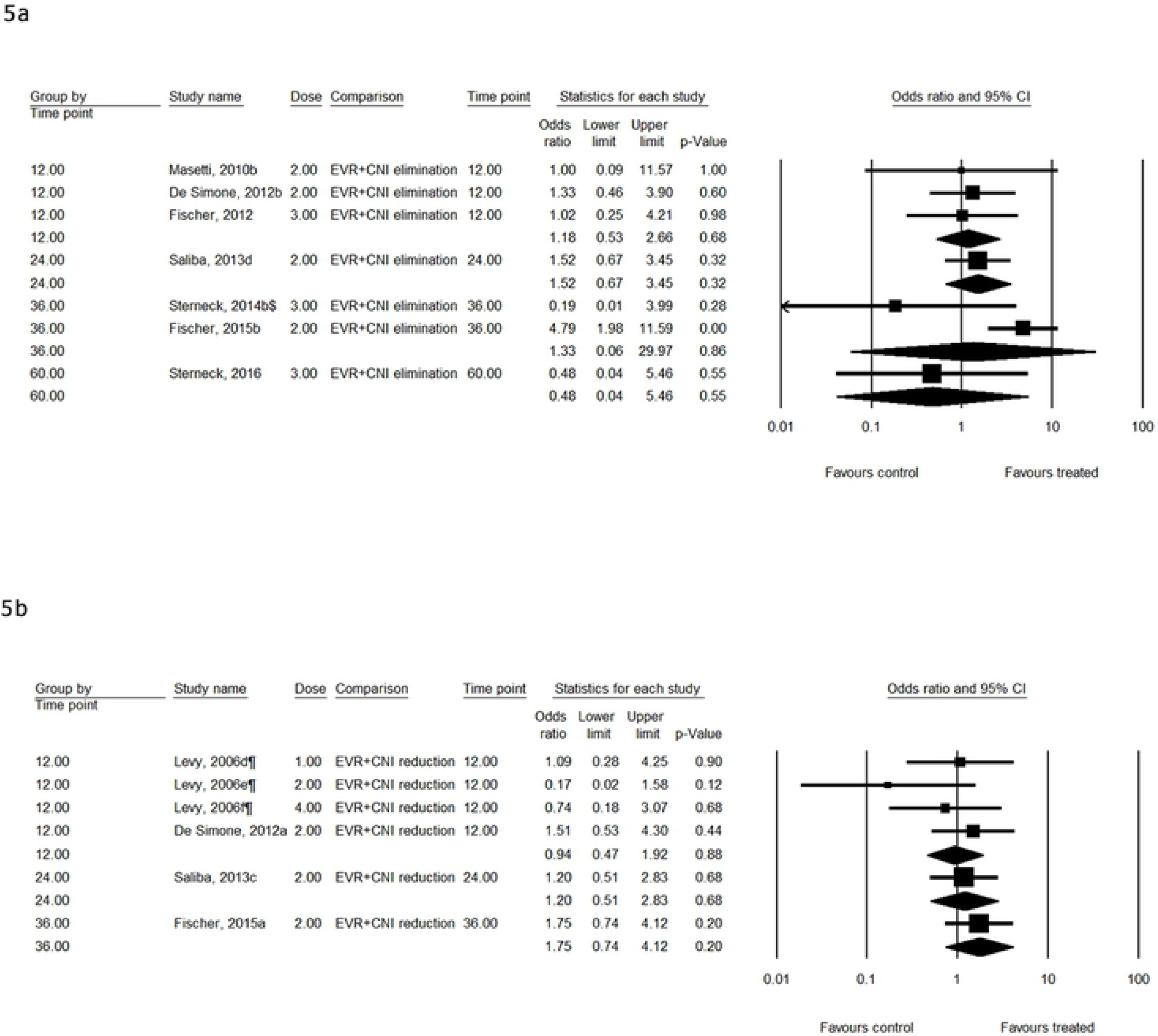

### Treatment emergent adverse events

Treatment emergent adverse events leading to treatment withdrawal were similar in treatment and control groups in EVR + CnI elimination trials at all the time-points. In EVR + CnI reduction trials, the odds of TEAE in treatment group were 2.37 (p=0.00) times higher at 12 months and 1.81 (p=0.01) times higher at 24 months as compared to the control group.

Subgroup analysis by dose showed that, in EVR + CnI elimination trials the odds of TEAE were similar in treated and control groups at 2 mg dose, whereas the odds of TEAE increased in treatment group at 3 mg dose (OR=1.95; p=0.03). In EVR + CnI reduction trials, the odds of TEAE were significantly higher in treatment group at 2 mg but not at 1 mg or 4 mg doses.

### Hepatic artery thrombosis

Incidence of hepatic artery stenosis was similar (P>0.05) between treatment and control groups of both therapy schedules, for all doses and at all-time points after liver transplant.

## DISCUSSION

This systematic review evaluates the recent evidence about the safety and efficacy of use of EVR in *de novo* liver transplant recipients. Meta-analysis was not possible due to insufficient RCTs with similar study design. A systematic review was therefore undertaken. The results of the review show that use of EVR either as monotherapy or in combination with reduced dose CnI (RTAC), can be beneficial in preserving renal function among patients undergoing liver transplantation. This is a result of CnI sparing, rather than a direct effect of EVR as the addition of EVR to standard CnI therapy having no effect on renal function^8^.

Important issues which need consideration in evaluating the results of this systematic review include the time of weaning of CnI therapy and the time of start of the EVR therapy; dose of EVR required for immunosuppression; reasons for discontinuation in the EVR groups; comparison of TAC elimination and RTAC regimes and the incidence of adverse effects with EVR as compared to CnI therapy.

Earlier trials had shown that late initiation of EVR after liver transplant, i.e. once renal impairment had developed, is not beneficial in decreasing the incidence of chronic renal failure^9,10^. In this review we therefore focused on studies involving *de novo* patients in whom EVR was started soon after transplantation, allowing early minimization or avoidance of CnI exposure. Both the PROTECT and H2304 studies have raised concerns about the time over which CnI therapy is reduced. Slow weaning (i.e. over 8 weeks) in the PROTECT study allowed the continuation of the CnI free (i.e. EVR) arm, whereas in the H2304 study, a similar treatment arm (TAC elimination) had to be discontinued because of clustering of episodes of BPAR around 120-180 days post randomization. The different protocols and discontinuation of the TAC elimination arm in H2304 preclude these two studies being analysed together and there is therefore only low quality evidence comparing TAC elimination with EVR+RTAC^13,16^.

The review shows that EVR+RTAC and EVR monotherapy were at least as effective as standard CnI therapy in preventing acute graft rejection and the composite efficacy end points^1,11^. However, use of EVR instead of CnI therapy by both Masetti et al ^10^ and Fischer et al ^1^ resulted in a decreased incidence of renal dysfunction. Most importantly, the progressive decrease in eGFR seen with standard CnI therapy was not seen in CnI sparing or CnI free regimens using EVR^11,12,13^, this difference achieved statistical significance at 36 months^13^.

One important concern highlighted by this review is the higher rate of treatment discontinuation in EVR containing regimens. The main reasons for discontinuation were proteinuria and infections. Proteinuria was the main adverse event leading to discontinuation of therapy during the initial two years, but was not seen in any patients from 24-36 months^13^. This might have been because of the limited number of patients who enrolled for the extension phase studies, but may also be due to a patient specific susceptibility that manifests within two years of exposure. Table 3 gives a comprehensive overview of the incidence of the most common AEs. In contrast, there was a decreased incidence of neoplasms in the EVR+RTAC arm in keeping with the known effects of mTOR inhibitors^13^. Levy et al suggested an increase in incidence of AEs with increasing dose, but this did not reach statistical significance^8^. They concluded also that the 4 mg/day dose may not be tolerated by liver transplant recipients. Further evaluation of the adverse events of EVR in liver transplantation is required.

This review is limited by the small number of RCTs identified, the difference in study design of the available RCTs, and the variable comparators in these studies. This was despite extensive search for RCTs, including both unpublished and published content. We had no language restriction, thus broadening our search.

In conclusion, the available RCTs showed that regimens containing EVR for *de novo* immunosuppression of liver transplant recipients allowing minimization of CnI exposure are at least as effective at preventing rejection and promoting graft survival as standard CnI therapy. Importantly, the studies identified demonstrated better renal function with EVR containing reduced CnI regimens as compared to standard CnI therapy. However, there is a need to evaluate the AEs with EVR regimens as compared to CnI therapy for both short and long term use. Everolimus therapy in combination with RTAC can be an alternative immunosuppressive therapy for liver transplant patients especially those with impaired renal function.

